# Extending and using anatomical vocabularies in the Stimulating Peripheral Activity to Relieve Conditions (SPARC) program

**DOI:** 10.1101/2021.11.15.467961

**Authors:** Monique C. Surles-Zeigler, Troy Sincomb, Thomas H. Gillespie, Bernard de Bono, Jacqueline Bresnahan, Gary M. Mawe, Jeffrey S. Grethe, Susan Tappan, Maci Heal, Maryann E. Martone

**Affiliations:** Department of Neuroscience, University of California San Diego, La Jolla, CA, United States; Whitby et al., Inc., P.O. BOX 90082 Indianapolis IN United States; Auckland Bioengineering Institute, University of Auckland, Auckland, New Zealand; Brain and Spinal Injury Center, University of California San Francisco, San Francisco, CA, United States; Department of Neurological Sciences, University of Vermont, Burlington, VT, United States; MBF Bioscience, Williston, VT, United States

**Author notes:** **Corresponding author:** Monique C. Surles-Zeigler. **Email Addresses and ORCID:** Troy Sincomb, Thomas H. Gillespie, Bernard de Bono, Jacqueline Bresnahan), Gary M. Mawe, Jeffrey S. Grethe, Susan Tappan, Maci Heal, Maryann E. Martone.

## Abstract

The Stimulating Peripheral Activity to Relieve Conditions (SPARC) program is a US National Institutes of Health-funded effort to improve our understanding of the neural circuitry of the autonomic nervous system in support of bioelectronic medicine. As part of this effort, the SPARC program is generating multi-species, multimodal data, models, simulations, and anatomical maps supported by a comprehensive knowledge base of autonomic circuitry. To facilitate the organization of and integration across multi-faceted SPARC data and models, SPARC is implementing the FAIR data principles to ensure that all SPARC products are findable, accessible, interoperable, and reusable. We are therefore annotating and describing all products with a common FAIR vocabulary. The SPARC Vocabulary is built from a set of community ontologies covering major domains relevant to SPARC, including anatomy, physiology, experimental techniques, and molecules. The SPARC Vocabulary is incorporated into tools researchers use to segment and annotate their data, facilitating the application of these ontologies for annotation of research data. However, since investigators perform deep annotations on experimental data, not all terms and relationships are available in community ontologies. We therefore implemented a term management and vocabulary extension pipeline where SPARC researchers may extend the SPARC Vocabulary using InterLex, an online vocabulary management system. To ensure the quality of contributed terms, we have set up a curated term request and review pipeline specifically for anatomical terms involving expert review. Accepted terms are added to the SPARC Vocabulary and, when appropriate, contributed back to community ontologies to enhance autonomic nervous system coverage. Here, we provide an overview of the SPARC Vocabulary, the infrastructure and process for implementing the term management and review pipeline. In an analysis of > 300 anatomical contributed terms, the majority represented composite terms that necessitated combining terms within and across existing ontologies. Although these terms are not good candidates for community ontologies, they can be linked to structures contained within these ontologies. We conclude that the term request pipeline serves as a useful adjunct to community ontologies for annotating experimental data and increases the FAIRness of SPARC data.

## Introduction

The Stimulating Peripheral Activity to Relieve Conditions (SPARC) program is a collaborative effort to document and describe the neural circuitry responsible for visceral control and to use this knowledge to promote the development of neuromodulation devices to improve organ function (National Institutes of Health, Office of Strategic Coordination-The Common Fund, 2021). SPARC comprises a consortium of researchers from multiple laboratories funded through multiple SPARC initiatives to identify autonomic nervous system (ANS) connectivity between end organs and the central nervous system. The effort also supports the production of tools and methods to understand this data.

The SPARC Data and Resource Center (DRC) is fielding infrastructure and tools for making these data and knowledge on ANS connectivity available to the research community and for use in models and simulations (Osanlouy et al., 2021). Data and tools are made available through the SPARC Portal https://sparc.science/. The SPARC DRC is charged with ensuring that all SPARC outputs, including data, knowledge about connectivity, models and simulations, adhere to the FAIR principles so that they are Findable, Accessible, Interoperable and Reusable (Wilkinson et al., 2016). Towards that end, SPARC outputs are curated to common standards, e.g., the SPARC Dataset Structure (Bandrowski et al., 2021), and are annotated to common semantic and spatial standards. To provide a common semantic underpinning to integrate across SPARC products, SPARC is utilizing community ontologies to annotate entities such as anatomical structures, organisms, and techniques. For spatial integration, SPARC is mapping experimental data on connectivity and molecular distributions onto common 2D maps and 3D organ scaffolds (Osanlouy et al., 2021).

As one of the main goals of the SPARC program is to generate detailed anatomical maps of the ANS, SPARC makes significant use of anatomical terminologies. Anatomical structures provide the common substrate across experimental data, computational models, 2D maps, 3D scaffolds, and a knowledge base of connectivity - the SPARC Connectivity Knowledge base of the Autonomic Nervous system (SCKAN), produced across the SPARC consortium. As such, it is critical to the SPARC program that anatomical terms are synced across efforts and that the necessary semantics are present to support queries and linkages across SPARC products. In support of FAIR principles, the overall strategy for the SPARC vocabulary is to utilize anatomical ontologies already in use by the community (Noy and Mc Guinness, 2001; Wilkinson et al., 2016). The SPARC vocabulary utilizes UBERON (Haendel et al., 2009; Mungall et al., 2012), the multi-species anatomy ontology, as the backbone of anatomical terminology efforts supplemented, as necessary, with additional species-specific ontologies such as Foundational Model of Anatomy (Nichols et al., 2014) and EMAPA (Hayamizu et al., 2013). The use of community anatomical ontologies also ensures that SPARC is interoperable with other programs in the Common Fund Data Ecosystem (The Common Fund Data Ecosystem | NIH Common Fund, 2021), a program and portal to allow cross query and integration of programs funded by the NIH Common Fund.

As SPARC is generating new data on ANS-end organ connectivity using advanced and varied techniques, investigators annotating data or modelers building detailed connectivity maps often require specialized terms that don’t appear in any community ontology (Balhoff et al., 2014). We, therefore, established a term submission pipeline using infrastructure initially developed by the Neuroscience Information Framework (Imam et al., 2012; Larson and Martone, 2013) for building and extending ontologies. Investigators and SPARC knowledge engineers may add new terms and relationships to the SPARC anatomical vocabularies through this pipeline, which subsequently become available for immediate use across SPARC. Where appropriate, terms are contributed to UBERON to enhance its coverage of the ANS. Given the key role anatomy plays in the organizing framework for SPARC products, we implemented a special review process for anatomical terms to ensure that they are clearly defined.

This paper provides an overview of the SPARC Vocabulary and describes the process and infrastructure for adding terms. We also provide an overview of the anatomical term review pipeline, analyzing the terms that have been submitted and reviewed to date. Finally, we discuss plans for further enhancement of the term request pipeline.

## Materials and Methods

### SPARC Vocabularies

The “SPARC Vocabulary” is a collection of terms and relationships used within the SPARC program for annotation, metadata, and search. The vast majority of these terms are derived from community ontologies in use across biomedicine (Figure 1). The SPARC Vocabulary uses the Neuroscience Information Framework Standard ontology (NIFSTD, https://github.com/SciCrunch/NIF-Ontology; RRID:SCR_005414) (Bug et al., 2008; Larson et al., 2009; Imam et al., 2011, 2012) developed by the Neuroscience Information Framework (RRID:SCR_002894). NIFSTD covers the major domains required for describing neuroscience data: Anatomy, Physiology, Molecules, Cells, Subcellular structures, Techniques, and Disease. NIFSTD itself is built through imports of major community ontologies and atlases, including the Uber-anatomy ontology (UBERON; RRID:SCR_010668) (Haendel et al., 2009; Mungall et al., 2012), the Allen Mouse Brain Atlas (RRID:SCR_002978; https://mouse.brain-map.org/) and other parcellation schemes. The SPARC Vocabulary also includes the Foundational Model of Anatomy (FMA; RRID:SCR_003379) (Nichols et al., 2014) for human anatomy and the Mouse Developmental Anatomy (EMAPA; http://www.obofoundry.org/ontology/emapa.html; RRID:SCR_021808) (Hayamizu et al., 2013) for mouse anatomy.

**Figure 1.**
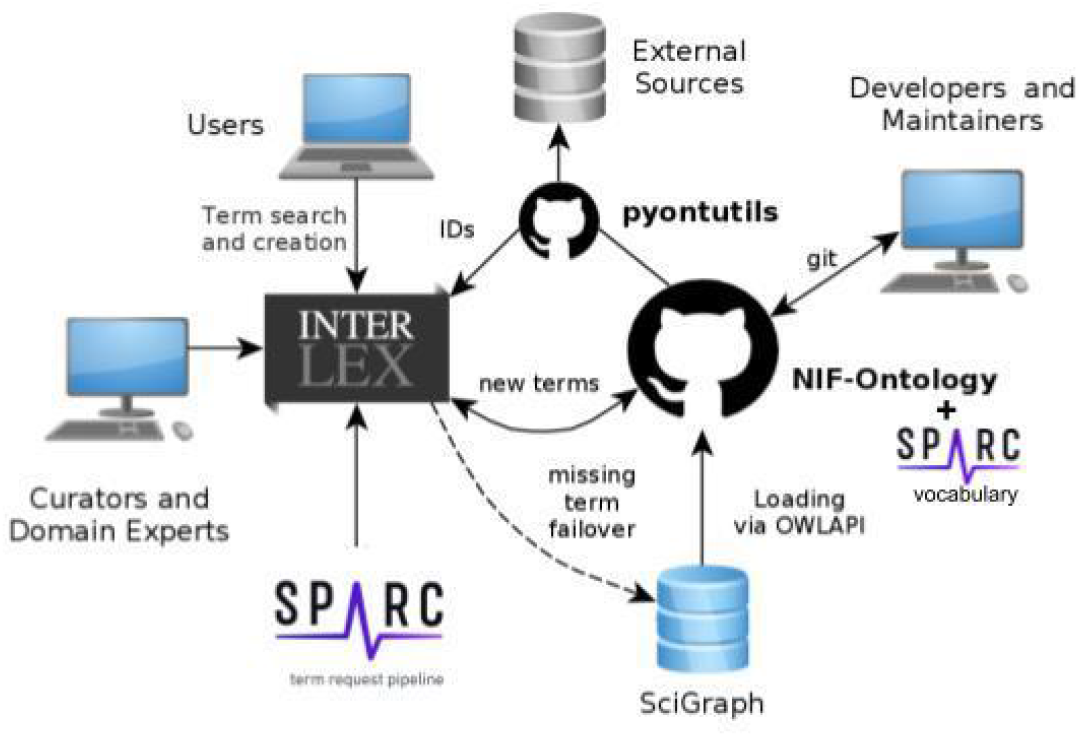
Schematic diagram of SciCrunch vocabulary infrastructure showing how content is managed across the different components. Different types of users interact with the system via different interfaces.

The SPARC vocabularies also contain terms and relationships that are contributed by SPARC investigators and developers. To add terms to the SPARC vocabularies and provide services to make these vocabularies available in SPARC tools. SPARC is utilizing the SciCrunch vocabulary management and services platform developed in part through the Neuroscience Information Framework, NIDDK Information Network (dkNET; RRID:SCR_001606), and the Center for Reproducible Neuroimaging Computation (ReproNim, https://repronim.org, RRID:SCR_016001) projects and maintained by the FAIR Data Informatics Lab (FDILab) at the University of California, San Diego (UCSD).

Access to the complete SPARC vocabularies in the form of a .ttl file, including complete ontologies imported by NIFSTD, is provided through GitHub (https://raw.githubusercontent.com/SciCrunch/sparc-curation/master/resources/scigraph/sparc-data.ttl). The process to merge this file is documented and automated here https://github.com/tgbugs/pyontutils/blob/master/nifstd/scigraph/README.org#sparc-sckan.

### SciCrunch Vocabulary Infrastructure

The SPARC vocabularies are served, accessed, and augmented through the SciCrunch vocabulary infrastructure shown in Figure 1. The SPARC vocabularies are housed in two primary stores:

1. Scigraph, a Neo4J-based graph database (https://github.com/SciGraph/SciGraph; RRID:SCR_017576) and
2. InterLex (https://scicrunch.org/scicrunch/interlex/dashboard, RRID:SCR_016178), an online vocabulary management system.

InterLex provides a user interface and workspace that allows search, viewing, addition, and editing terms and relationships. These two components are described in more detail below.

#### SciGraph

Scigraph is a Neo4j graph-based OWL ontology database that serves the reasoned version of an ontology so that it can be used in information systems. SciGraph replaced the original Ontoquest database developed by NIF used for their semantic search function (Gupta et al., 2008). SciGraph can be queried through Cypher queries to traverse the relationships in the graph. Within SPARC, SciGraph is used as an ontology lookup service that accesses the NoSQL Neo4j graph for the SPARC vocabulary through a REST API. The programmatic way to view and search for terms within the SPARC vocabulary is through the Ontquery python package (https://pypi.org/project/ontquery/; RRID:SCR_021659) initially developed by ReproNim and the BRAIN Initiative Cell Census Network, BRAIN Initiative’s Cell Census Network (BICCN, https://biccn.org/, RRID:SCR_015820) Brain Cell Data Center (BCDC).

#### InterLex

InterLex is a web-based vocabulary management system with a user interface (https://interlex.org; RRID:SCR_016178) and a database of medical and biological terms. InterLex replaced NeuroLex (Larson and Martone, 2013), an online semantic wiki for viewing and extending the NIFSTD ontologies, and contains all NeuroLex terms. Additional terms were added to InterLex through bulk uploads from external ontologies such as Mondo, Uberon, and FMA. This upload is done via a semi-manual process of merging terms with Internationalized Resource Identifiers (IRI) mappings provided by external ontologies and curating remaining terms based on their labels, annotations, and relationship properties. This foundation of merged ontologies is designed to allow a non-expert to search, view, and add terms and define relationships between them. Each new term is given a full Uniform Resource Identifier (URI) with a unique ilx: prefix, immediately referenced. InterLex also maps terms to URIs from external ontologies. Each term can specify the default identifier to be used with that term (e.g., the UBERON identifier should be used instead of the InterLex identifiers).

InterLex data is held within a MariaDB (https://mariadb.org/) relational database (Forta, 2011; Bartholomew, 2012), and the underlying schema represents terms, relationships between terms, and term annotations using a Resource Description Framework (RDF) model. InterLex’s internal identifiers support automatic versioning by using a simple document versioning pattern where the current term versions are stored in one database table, and all previous versions of the terms are stored in another table. A subset of the contents of these MariaDB tables are internally transformed into a single turtle file (.ttl) and loaded into SciGraph after being added to NIFSTD as the sparc-community-terms.ttl file found here (https://raw.githubusercontent.com/SciCrunch/NIF-Ontology/sparc/ttl/sparc-community-terms.ttl). The InterLex database automatically syncs with an Elasticsearch index (Divya and Goyal, 2013) to enable term search via a simple REST API or through the InterLex interface. Access to the REST API requires an API Key, retrievable with a SAWG community account created at https://scicrunch.org/sawg/join. In addition, the Python API wrapper, Ontquery also incorporates the InterLex REST API to streamline pipelines for external users. This feature provides an ontological foundation for the scientific community to use the backend of InterLex to search, add, modify, and comment on term entities.

The InterLex user interface (https://scicrunch.org/scicrunch/interlex/dashboard, RRID:SCR_016178) is housed in a research portal within the collaborative SciCrunch Infrastructure (Whetzel et al., 2015). The SciCrunch framework provides a platform and associated tools for the creation of community data portals on top of a common set of resources. SciCrunch currently supports multiple public and private community portals within this shared infrastructure (public portals are listed at: https://scicrunch.org/browse/communities). InterLex is a component of the SciCrunch platform, allowing communities the ability to work with and view terms contributed via a specific community. However, all terms contributed by a community are immediately available to all InterLex users. To support the SPARC program, we established a custom portal for the SPARC Anatomy Working Group (SAWG) available at https://scicrunch.org/SAWG. When terms are added to InterLex through a community portal, the terms are automatically tagged as entering via SAWG and may be viewed by a custom community dashboard. All terms contributed via the SAWG portal are available at https://scicrunch.org/sawg/interlex/dashboard-history?origCid=504&page=1&sort=desc.

As of August 2021, InterLex contains over 400,000 terms, 55 relationship types, 59 annotation types shared across 9 communities. A full set of sources imported by InterLex is given on the InterLex home page (https://interlex.org). In addition to vocabularies like MeSH and ontologies such as UBERON, InterLex also has imported NIH Common Data Elements and other term sets. These sources provide terms for InterLex, but the relationships between terms are usually not fully preserved. InterLex is not meant to be a comprehensive source for these ontologies; rather, terms from these ontologies are imported on an as-needed basis to support the knowledge engineering required for linking across terms, as described in a later section. Accordingly, InterLex contains only a subset of the SPARC Vocabulary, best characterized as the subset of SPARC Vocabulary, that is currently used in SPARC across data sets, models, scaffolds, and knowledge bases like SCKAN. The relationship between the SPARC Vocabulary and the infrastructure components is shown in Fig 2.

**Figure 2.**
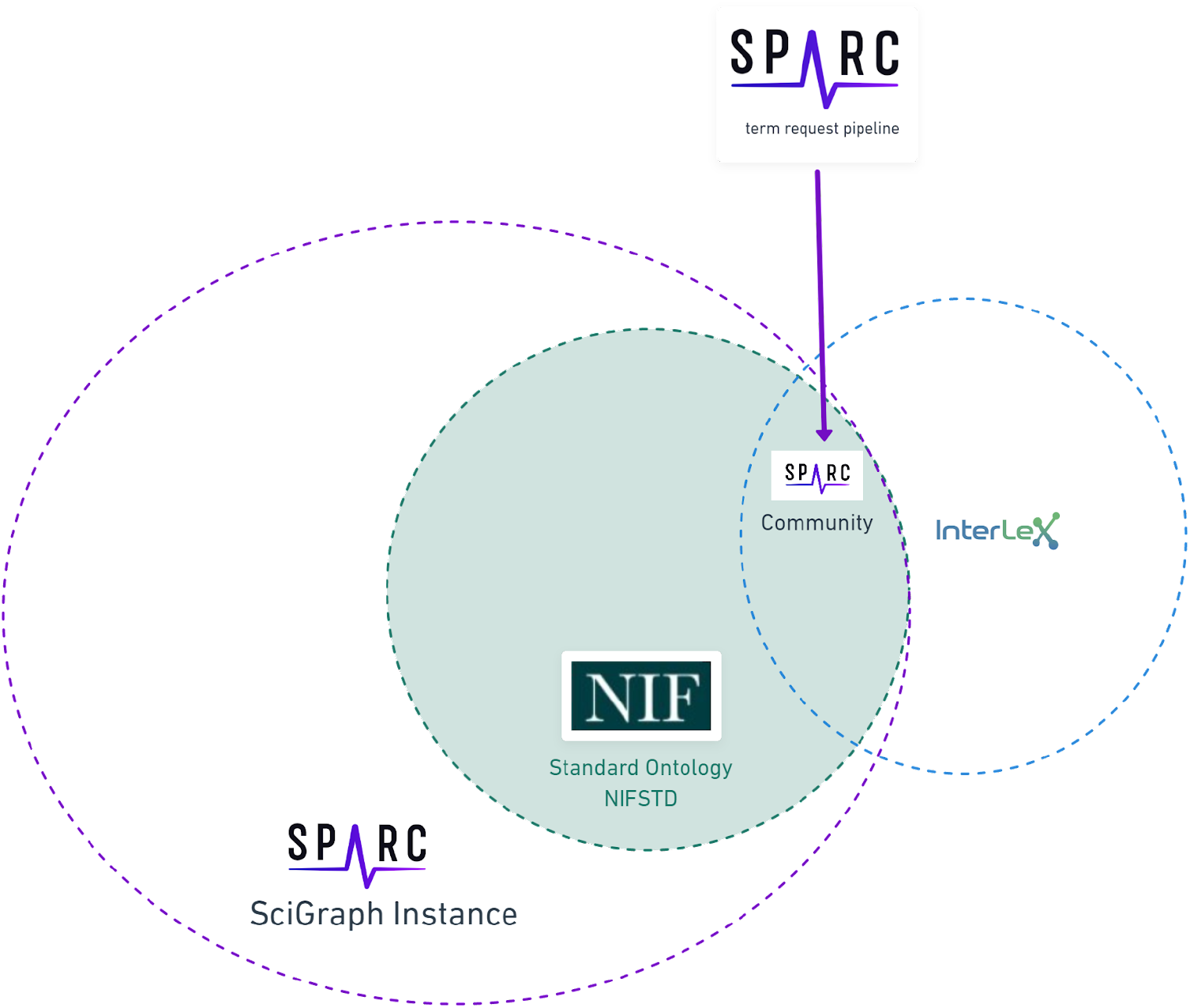
The relationship of the SPARC vocabulary to infrastructure components. The SPARC Vocabulary is represented by the large circle on the left (solid dotted line). The entire SPARC Vocabulary is available through a SciGraph instance, including the full imports of community ontologies comprising the vocabulary. The totality of InterLex is represented by the small circle on the right (blue dotted line) that partially exists within the SciGraph instance. The overlap between these circles represents the subset of the SPARC Vocabulary that is available in both InterLex and SciGraph that is made available via the SPARC Community Portal (purple dotted line) and augmented through the term request pipeline.

### Use of SPARC vocabulary within SPARC

The SPARC Vocabulary provides the common semantic framework for integrating and querying across SPARC data sets, models, maps, simulations, and spatial coordinate systems. Collectively, we will refer to these as SPARC products. In the following, we describe some of the main usage scenarios.

#### SPARC Datasets

SPARC investigators submit data to the SPARC data platform, Pennsieve (previously Blackfynn, https://app.pennsieve.io/; RRID:SCR_021677), where it undergoes human and semi-automated curation to the SPARC Data Set Structure and Minimal Information Standard (Bandrowski et al., 2021; Osanlouy et al., 2021). Each data set must be accompanied by a detailed experimental protocol deposited in Protocols.io. Metadata provided by the investigator is mapped to the SPARC vocabularies by a human curator supported by semi-automated mapping tools. The complete SPARC dataset submission, curation, and registration pipeline are illustrated in Figure 3. The points where vocabularies are applied for annotation are indicated by stars.

**Figure 3.**
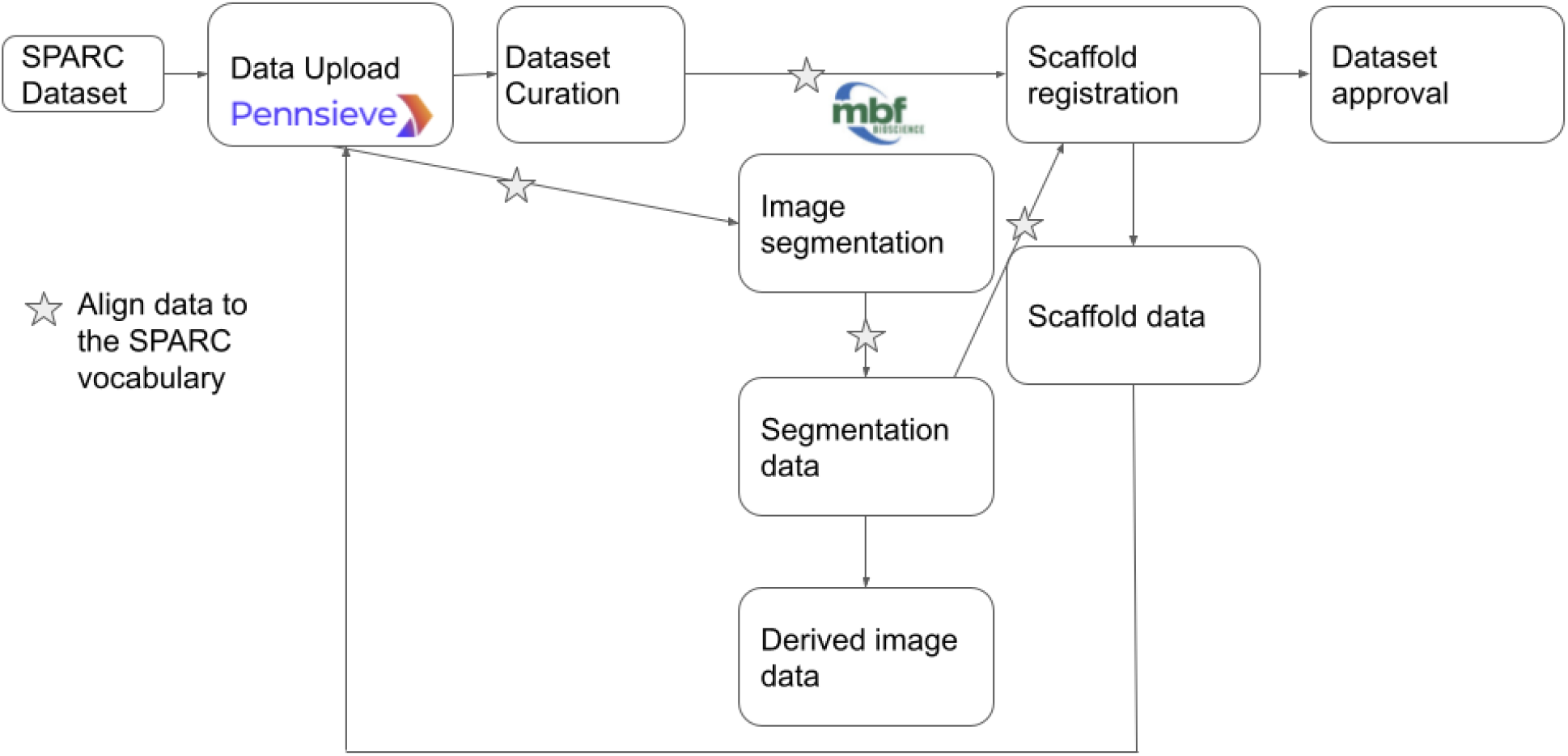
Overview of SPARC data workflow modified from (Osanlouy et al., 2021). Stars indicate steps in the pipeline where the annotation is performed, and terms are most likely to be added to the SPARC Vocabulary.

#### Microscopy Images and Scaffolds

SPARC investigators acquire detailed and diverse 2D and 3D microscopy images and mappings of cells, anatomical structures, projections, and molecules in the ANS. To aid in 3D reconstruction, segmentation, and annotation of these images, many researchers employ neural reconstruction and anatomical mapping software from MBF Bioscience (RRID:SCR_004314), such as Neurolucida 360™ (RRID:SCR_016788) and Tissue Mapper™(RRID:SCR_017321). In order to compare distributions of cells and projections within and across datasets, these distributions are spatially registered to one or more 3D computational scaffolds of major organs (Osanlouy et al., 2021) by SPARC members at the Auckland Bioengineering Institute (ABI). Fitting scaffold models to the experimental data is done by identifying a common set of fiducial points within these images and demarcating concordant points in the segmentation data and the scaffold model. The segmentation data, stored according to MBF Bioscience’s XML neuromorphological file specification (Angstman et al., 2020; Sullivan et al., 2021), is then ingested into ABI’s ScaffoldFitter software (RRID:SCR_019002). Each dataset is registered to an individual scaffold, transforming the data into a common coordinate space, making it possible to integrate or average multiple datasets in this space.

This process is made possible due to a special integration within MBF software that communicates with a SciGraph database instance through an API, allowing one to tag segmented anatomical structures and fiducial points to the SPARC vocabulary (Figure 4). From MBF software, a user can select a button “Vocabulary Services” to get instant access to the up-to-date term lists within the SPARC vocabulary. The user is first prompted to add additional metadata about their subject and is given options via a series of drop-down menus to select terms sets related to specific organs, species, or atlases (parcellations). After the selection is made, the tool provides a list of all terms associated with that organ system that can be used to name or classify segmented neural structures, vasculature, anatomies, and whole cells from their microscopy image data. Terms can be found by manually searching the list provided or by using an auto-complete function. Although all investigators are encouraged to use terms within the SPARC vocabulary to annotate their images, they may enter custom terms if needed. Custom terms can describe structures or act as a placeholder until that term (or term set) is requested. These terms serve as a conduit to the Anatomical Term Request Pipeline (see the section below), which is accessible from MBF software to facilitate this process (Figure 4D).

**Figure 4.**
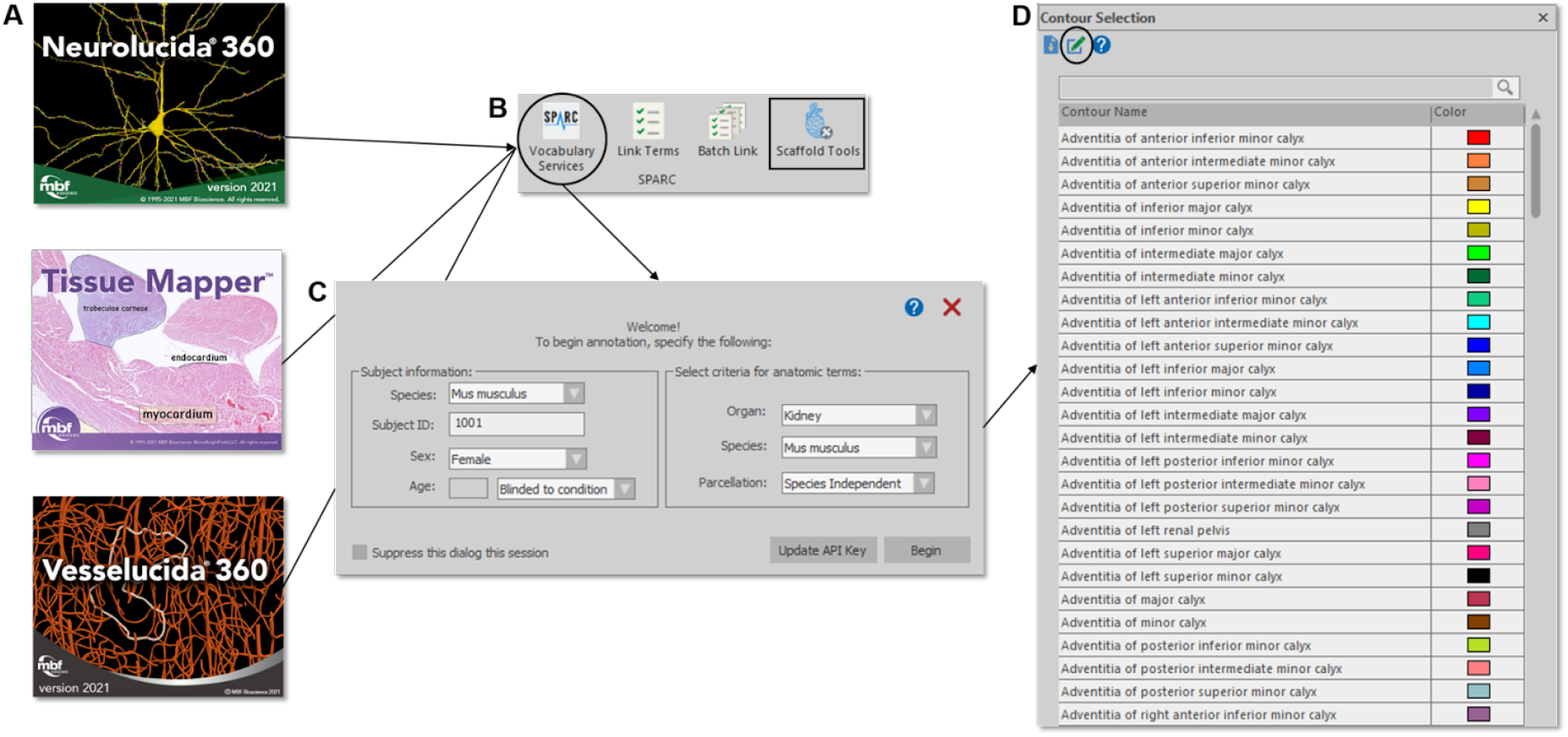
Accessing the SPARC Vocabulary through an API in the MBF Bioscience software suite (A). The Vocabulary Services tool (B, circle) provides users with a subset of SPARC Terms (D) – specified by organ, species, and parcellation (C) – that can be applied to anatomical annotations. An option to request new terms (D, circle) launches the SAWG portal dashboard. SPARC Terms are also available through Scaffold Tools (B, square), where segmentation data and organ scaffolds are displayed side-by-side to preview concordant fiducial SPARC terms for subsequent scaffold registration.

With the understanding that data is often generated prior to the full maturation of software tools and vocabularies, MBF added a software feature that permits investigators to revisit segmentation data files to programmatically and comprehensively (with batch functions) add SPARC vocabularies and their associated metadata to XML files at a later date (Figure 4B).

At a file level, MBF’s neuromorphological file format stores the globally unique and persistent ontological identifiers as properties (e.g. *<contour name= “*Vagus nerve”…*<property name=“TraceAssociation”><s>*http://purl.org/sig/ont/fma/fma5731*</s>*…) that are machine-readable for increased interoperability with other tools (e.g. ScaffoldFitter) and searchable on open data portals (e.g. sparc.science).

#### SPARC connectivity models

Investigators within SPARC are producing detailed models of ANS connectivity, which include the granular routes via which neurons travel within the body using the ApiNATOMY platform (de Bono and Hunter, 2012; de Bono et al., 2014, 2018; Kokash and de Bono, 2021). ApiNATOMY is a knowledge model for biological connectivity and includes a set of tools that create anatomy schematics overlaid with ontological information. These tools are used to build and annotate circuit graphs, i.e., wiring diagrams of the peripheral nervous system processes. The example shown in Figure 5 is a diagram representing the neuronal connections between the spinal cord and the bladder to consolidate and query knowledge about urinary system innervation (produced by Surles-Zeigler et al., 2021).

**Figure 5.**
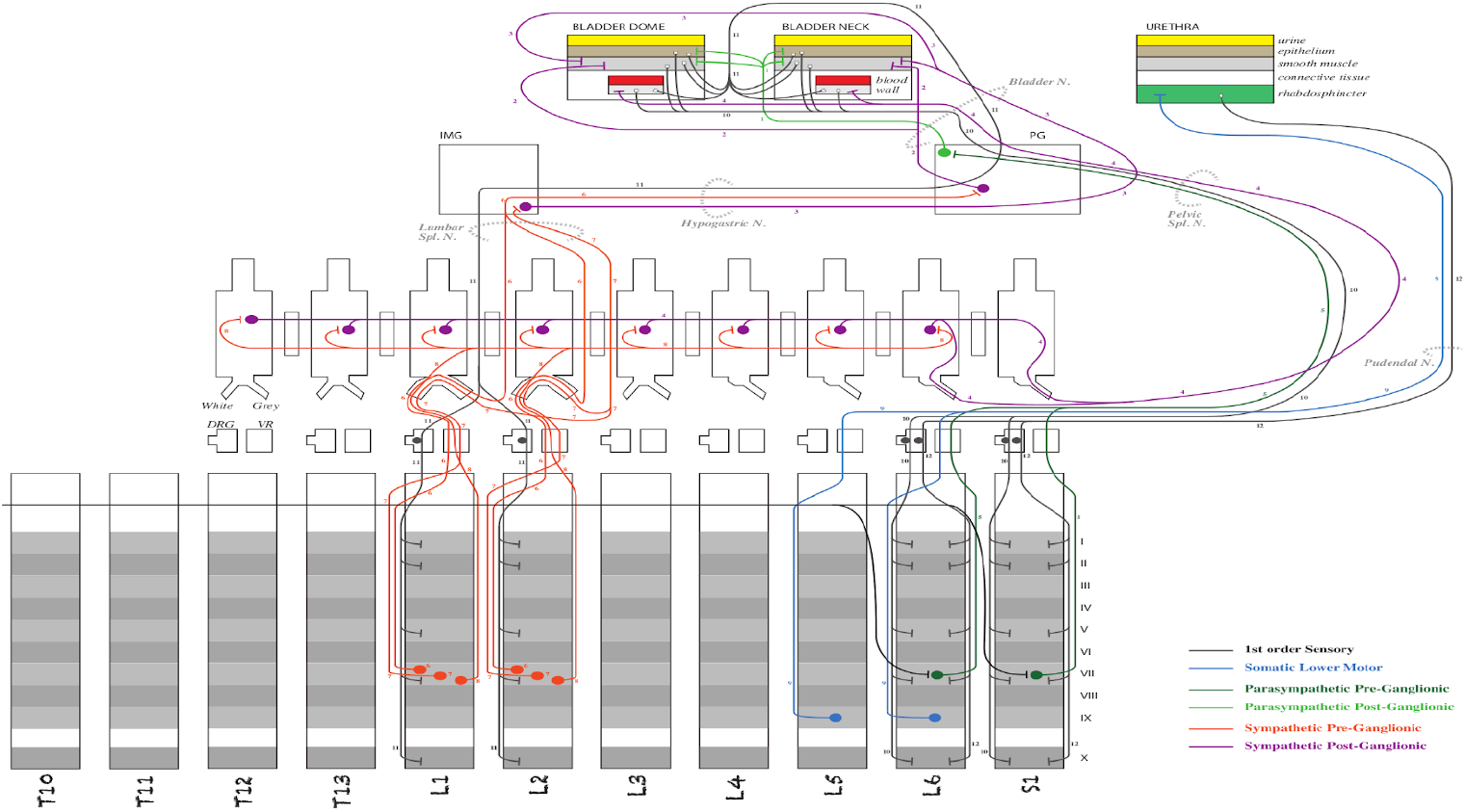
Schematic diagram of the ApiNATOMY model of bladder innervation (Surles-Zeigler et al., 2021). The diagram illustrated major neural circuits involving the urinary bladder and urethra.

### Anatomical Term Request Pipeline

Once someone with a SAWG Portal account adds a term to InterLex, it becomes immediately available for annotation. However, because of the central importance of anatomy across all SPARC products, anatomical terms must go through specialized pipelines and a curatorial review before being added to the SPARC Vocabulary. The SPARC anatomical term request pipeline is an iterative process that consists of three main steps: term request, term review, and term engineering (Figure 6). Each of these steps is described in more detail below.

**Figure 6.**
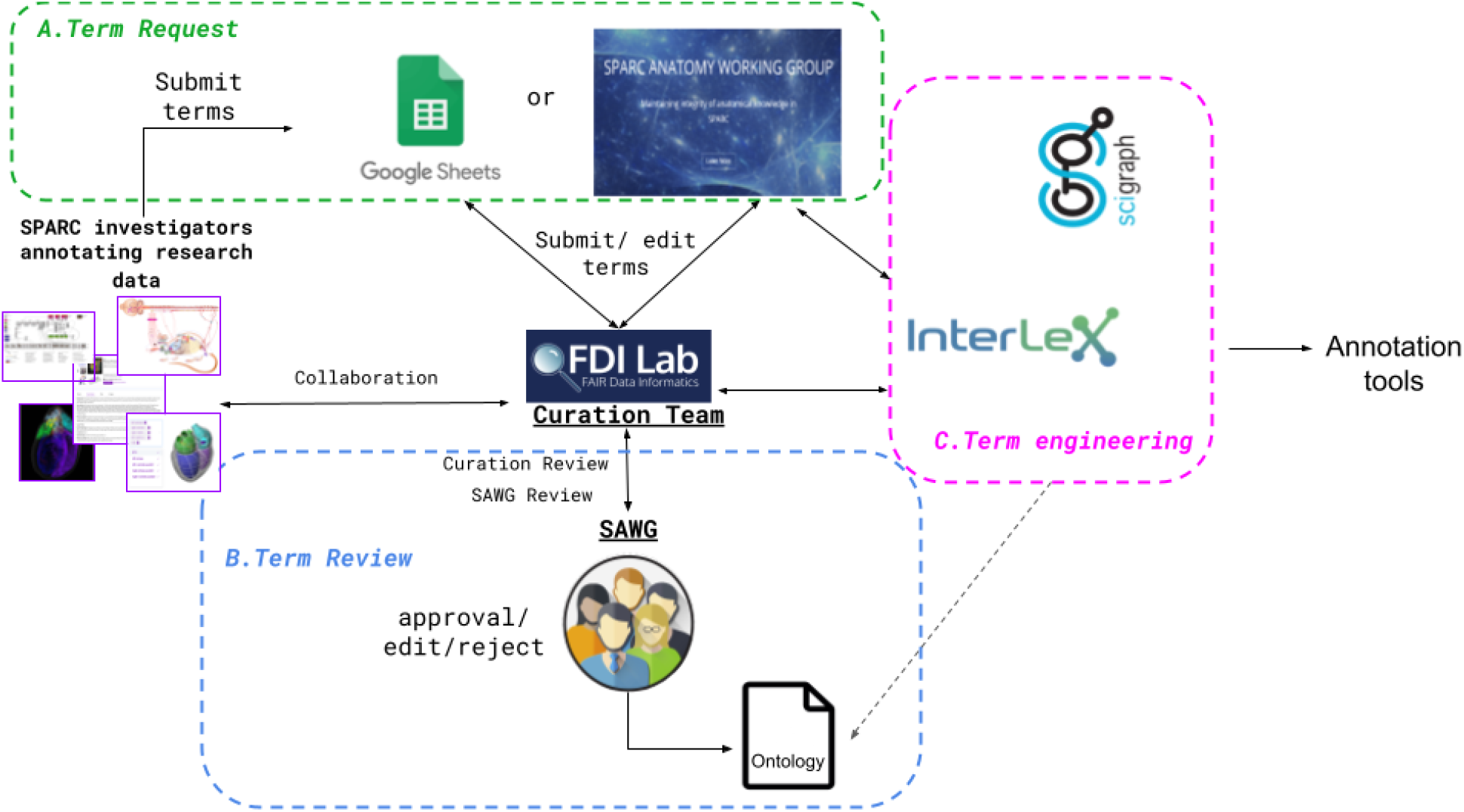
SPARC term request and review pipeline. The process comprises three iterative steps: A) submission (Term request), B) Term review, and C) Term engineering, shown in the 3 colored boxes. Details are provided in the text.

#### Term Request

Term(s) are submitted to the term request pipeline through the SAWG dedicated portal within SciCrunch (https://scicrunch.org/sawg). Single terms are usually added directly into InterLex, where they are tracked through the SAWG portal dashboard: https://scicrunch.org/sawg/interlex/dashboard-history?origCid=504&page=1&sort=desc. The portal dashboard is checked weekly by curators for any new terms. As an account is needed to add terms to InterLex, the terms are all identified by the submitter’s name and email address. They are also automatically tagged to the SAWG community.

When multiple terms are requested, they are generally submitted via the SPARC Term Request Google Sheet (RRID:SCR_017679) accessible via the SAWG portal. Terms requested via the MBF tools use this route. Terms submitted via the Google Sheet are accompanied by additional metadata such as the requestor’s email and name, date of submission, the investigator that contributed the term, definition, definition source, and any additional notes about the term. The SPARC term curation team at UCSD receives a notification for each change made to the sheet and can communicate with the requestor via comments within the sheet.

#### Term Review

All anatomical terms submitted to the SPARC vocabularies undergo additional curation. The entire review process is presented as a decision tree in Figure 7. The initial review is completed by the UCSD SPARC curators, who determine whether the term already exists within the SPARC vocabulary or is a known synonym of an existing term. If it does exist, the team reviews the metadata and relationships available with the term to ensure they are complete and reflect the intended usage by the author. For example, if a term enters the term request pipeline through the Google sheet and the curator identifies it in InterLex, the curator may still add a definition, even if the term itself is not added. If the term is not present in the SPARC Vocabulary, the curator does a review to determine whether it may be present in another community ontology by searching InterLex and BioPortal (https://bioportal.bioontology.org/). If so, it is entered into the SPARC vocabulary via InterLex and cross-mapped to any external identifiers.

**Figure 7.**
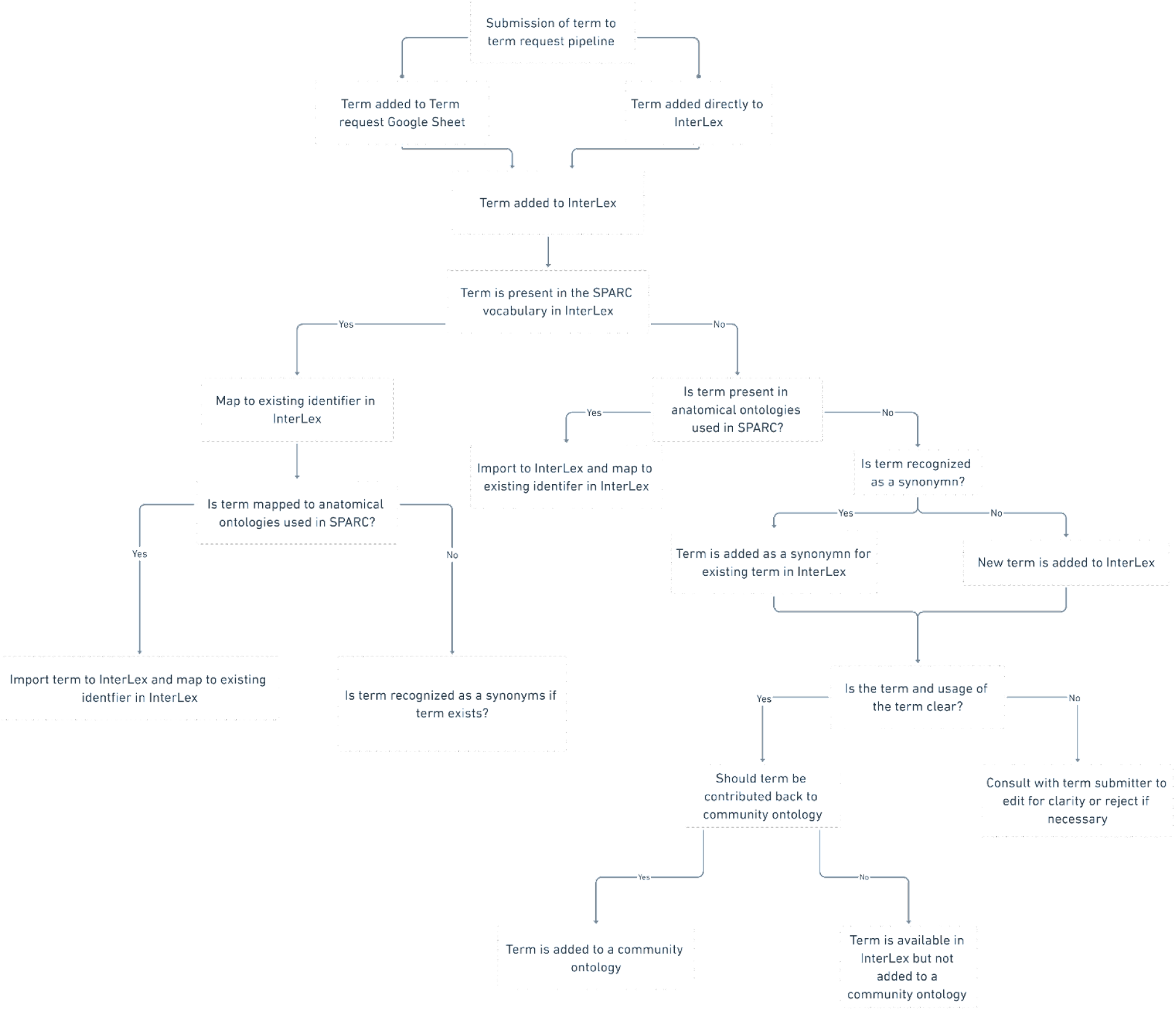
SPARC term request workflow. Flow chart illustrating the term request and review steps of the term request pipeline

After initial review, the terms are presented to the SAWG weekly for review. The SAWG provides independent expertise and arbitration on the anatomical terms submitted to the pipeline. Terms are reviewed to determine whether the label is correctly formulated, the definition is clear, and the term is recognized as an anatomical term. If the members of the SAWG do not have the required anatomical expertise to adjudicate the term, they do additional research or consult outside expertise. The SAWG may request further clarifications or information from the submitter when the use of the term is unclear, e.g., an annotated image. The SPARC curator may also consult the SAWG regarding mapping a submitted term to terms within the vocabulary, e.g. if the term is a synonym or child of an existing term.

Terms reviewed by the SAWG are (1) approved, (2) revised, or (3) rejected. If a term is approved, a message is sent to the requestor confirming that the approved term is present and will be available in all instances of the SPARC vocabularies at the next release (including SciGraph). The metadata in InterLex is updated to include the annotation property “ApprovedBySAWG’’ to note the SAWG has approved the term. If a revision is recommended, the investigator or point of contact is contacted to approve the edit. The edited term is then changed within InterLex. Lastly, a term can be rejected if it is thought to be erroneous in some way. If the term is rejected by the SAWG, the investigator or point of contact is contacted to provide more information or asked to use an alternative term(s).

At the recommendation of the SAWG, terms representing general anatomical structures are contributed back to community ontologies to enhance their coverage of central nervous system-peripheral nervous system-organ interactions. If a term is accepted by a community ontology, its identifier is entered as the preferred identifier and is mapped to the InterLex identifier.

#### Term Engineering

The term engineering is a critical step and occurs at multiple stages in the pipeline. During this step, the terms within InterLex are given formal definitions instantiated in a set of relationships, that link them to other terms in a manner that serve SPARC use cases. The curation team performs term engineering. It includes ensuring that terms are linked to identifiers in outside ontologies where appropriate, are classified under the correct parent term through the “is a” relationship and other structures through “part of” relationships.

Other relationships and annotations may also be added to ensure that submitted terms are linked to community ontology terms. One example of this are terms supplied by the MBF mapping pipeline described in the Materials and Methods subsection, “Microscopy Images and Scaffolds”. Anatomical terms may be missing from the organ-specific list in the MBF software, although present in the SPARC vocabulary. In these cases, we provide a shortcut relationship to bind these terms to the appropriate organ list through the “includeForSPARC” relationship making the terms findable in these lists. This relationship ensures that the term is tied to the appropriate organ terms in UBERON or FMA via SciGraph without implying that these relationships are sanctioned by these source ontologies.

InterLex also allows for the insertion of annotation properties which enables additional provenance or tags to be added to a term. Tags provide efficient traversal while also reducing runtime of dynamically pulling complete datasets, including datasets pertaining to SPARC. A complete set of relationships including annotation properties currently in InterLex can be found at: https://scicrunch.org/scicrunch/interlex/search?types=relationship,annotation

## Results

The SPARC Term Request Pipeline has been operational since October 2019. From October 2019 through August 2021, there have been 312 anatomical terms submitted to the pipeline identified by SPARC members as missing in the SPARC vocabulary. Users of the SPARC Vocabulary have several options for access: 1) Programmatic access to the full ontology via SciGraph, which requires the use of an API and familiarity with the Cypher language; 2) Programmatic access to the subset of the vocabulary in InterLex via Elasticsearch based APIs; 3) Access to the subset of the vocabulary in InterLex via a user portal; 4) Access to the SPARC organ-specific term sets via the MBF tools. When a term is labeled as “missing”, it can therefore mean missing from the SciGraph instance, missing from InterLex, or missing from the organ-specific term list, depending on how the vocabulary was accessed.

Of the terms submitted to the pipeline, 161 terms were submitted through the SPARC term request Google Sheet, and 151 terms were added directly into InterLex. The majority of the terms were requested by curators at MBF Bioscience (129 terms) to assist SPARC investigators with annotating data within their software. The remaining terms were used to annotate ApiNATOMY models (124 terms), 3D scaffolds (44 terms), and SPARC datasets (5 terms). Ten terms were added by curators to connect submitted terms to existing terms. The disposition of terms currently in the pipeline is shown schematically in Figure 8.

**Figure 8.**
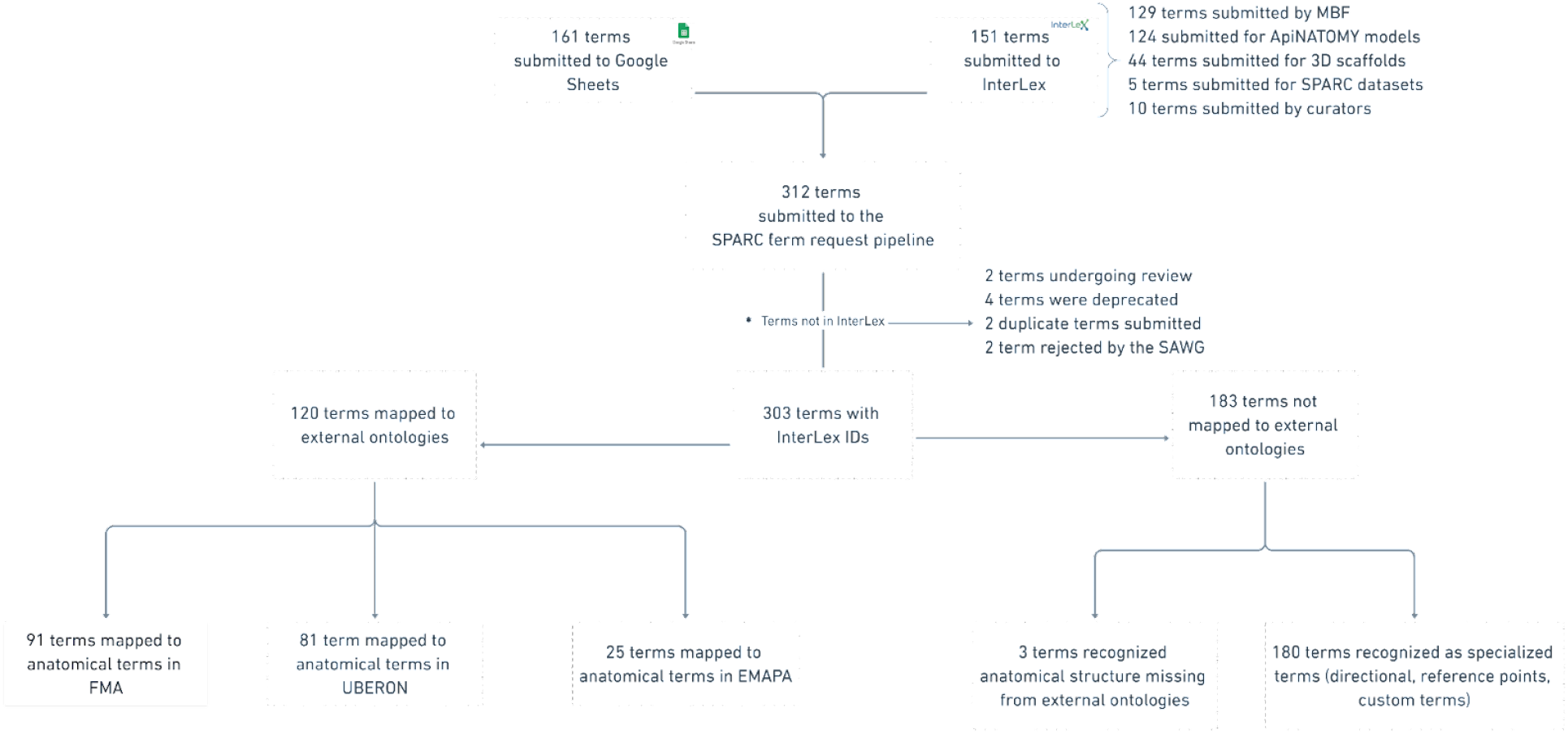
Disposition of all terms submitted to the term request and review pipeline as of August 2021. The star in the figure notates the 10 terms labeled as “not in InterLex ‘‘. This label refers to terms that were added to the pipeline but were either not added to the SAWG community or depreciated from InterLex, a subset (SAWG community) of the vocabulary in InterLex via free-text search.

Sometimes, a term request highlighted a large gap in the SPARC Vocabularies that could be filled by importing a new ontology. For example, term requests for mouse-specific anatomy led to the import of the EMAPA ontology, significantly expanding the size of the SPARC Vocabulary. However, here we focus only on the actual terms requested.

### Curation term review

Of the 313 anatomical terms entering the pipeline,10 terms were not added to the SPARC community in InterLex: 2 were rejected by the SAWG, 6 terms were found to be duplicates and 2 terms are currently under review. Of the duplicate terms, 2 were submitted via the Google Sheet and the curator determined they already exist in the SPARC Vocabulary, and 4 terms added to InterLex were deprecated after they were found to be duplicates. The 4 deprecated terms were redirected to the original term in InterLex with the “replacedBy’’ relation tag. In total, 303 anatomical terms were added to the SAWG community within InterLex.

Of the 303 terms in the pipeline, 120 terms (40%) already existed in community ontologies, FMA (n= 91), UBERON (n=81), EMAPA (n=25). Almost half of the terms had mappings to both FMA and UBERON (74 terms). Many of these terms were common anatomical structures, e.g., lung (UBERON:0002048, FMA:7195), brain (UBERON:0000955, FMA:50801), colon (UBERON:0001155, FMA:14543), and solitary nucleus (UBERON:0009050, FMA:256691), but were not found when needed.

The majority of submitted terms (N =183, 60%) did not directly map to community ontologies. These non-mapped terms were primarily of two types: (1) recognized anatomical structures that were missing from community anatomical ontologies, and (2) custom terms used within SPARC for registration of SPARC data or analysis. As part of the knowledge engineering step, all submitted terms added to the community were connected to a community ontology through one or more relationships, when possible. The major relationships used for term engineering of the 313 terms are shown in Table 1.

**Table 1.** Relationships entities used within the term request pipeline

| Relationship | InterLex ID | Definition | count |
| --- | --- | --- | --- |
| Is part of | ILX:0112785 | Generic partonomy relationship; does not distinguish among subtypes | 94 |
| replacedBy | ILX:0383242 | Use on obsolete terms, relating the term to another term that can be used as a substitute | 4 |
| includeForSPARC | ILX:0738400 | A relationship that binds a term to the required entities for the purposes required by SPARC, e.g., returning a term in response to a query for all relevant organ parts when it is not specified in the core ontology. We view this as a temporary and practical solution. At some points, all such terms will be contributed back to the core ontologies for proper engineering. | 208 |
| intersectionOfPartOf | ILX:0739238 | The relationship that exists between 2 given classes (sets) and contains only an arbitrary part of the elements common to both classes. This is both a logical and spatial intersection. When presented as an anatomical entity, this relationship is treated as an intersection of its parts. | 55 |
| marksAnatomicalJunctionOf | ILX:0739265 | A fiducial marker. It marks an anatomical reference point in an image between two or more given anatomical regions (classes). | 30 |
| anatomicalJunctionOf | ILX:0739272 | A relationship between an anatomical junction and a part. | 6 |
| hasDefinitionSource | ILX:0739292 | Source of definition if source another ontology | 38 |
| includesTerm | ILX:0770273 | Includes entity of type term. Used to create term sets. | 94 |

In the first group, 3 terms were determined by the SAWG to be bona fide anatomical structures that should be contributed to UBERON: Anterior subdiaphragmatic vagus nerve (ILX:0738436), Inner submucosal nerve plexus (ILX:0777077), and Outer submucosal nerve plexus (ILX:0777078). The second group (177 terms) consisted of specialized terms that were used for annotation, segmentation, or registration of SPARC data. This group included: (a) directional terms, (b) terms that describe a reference point between multiple anatomical structures, and (c) custom anatomical terms that are not in common use. In all 3 cases, these terms could not be directly mapped to a single existing ontology class but could be constructed with combinations of existing terms and relationships. For example, the term “junction between pulmonary valve and right ventricle” (ILX:0777101) is composed of: “pulmonary valve” (UBERON:0002146, FMA:7246), “heart right ventricle” (UBERON:0002080, FMA:7098) and the relationship “ “marksAnatomicalJunctionOf” (ILX:0739265).

Directional terms comprised 55 terms that contained relative directional qualifiers within the term, e.g., dorsal or posterior, but were not recognized as bona fide structures in common parcellation schemes or the scientific literature. An example of this type of term is “Dorsal part of urinary bladder lumen” In this case, urinary bladder lumen is an anatomical structure that already exists within an ontology, while the term “dorsal part” is used as a descriptor. Within InterLex, these terms were related to their parent structure using the predicate “intersectionOfPartOf” as illustrated in Figure 9.

**Figure 9.**
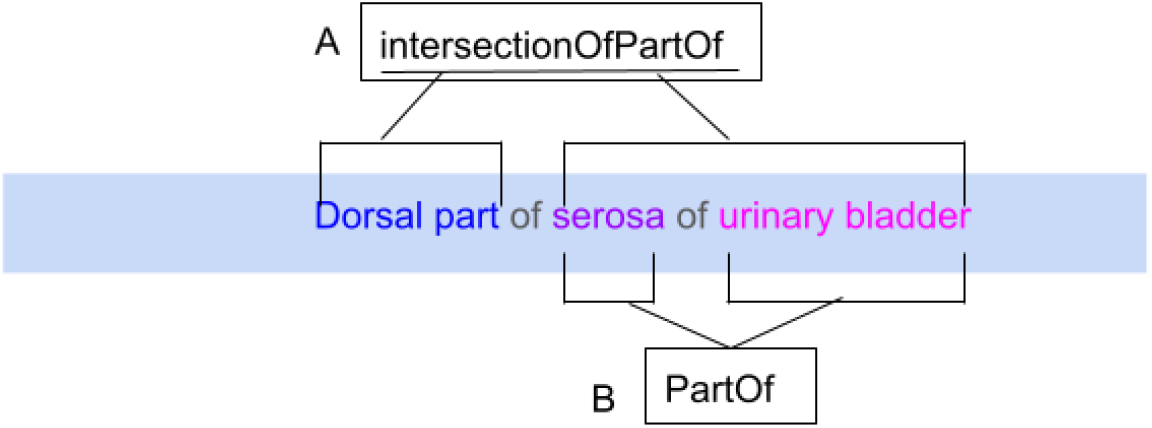
Term Engineering for the submitted term “Dorsal part of serosa of urinary bladder”. The term engineering is usually performed by adding appropriate machine-readable relationships to relate submitted terms to the appropriate supercategory and other terms. (a) An example of a relationship term in InterLex is IntersectionOfPartOf. (b)This relationship acts as a linker between the term “Dorsal part” with identifier PATO:0001233 and “serosa of urinary bladder” with identifier UBERON:0001260.

Reference point terms comprised 16 terms classified as fiducial markers or a point of reference while annotating data for registration to a 3D organ scaffold. An example of a term in this sub-group is “Junction between inferior cardiac nerve and cervicothoracic ganglion” (ILX:0739267), where this junction is used to notate a point on a scaffold between the origin of the inferior cardiac nerve as it intersects with the cervicothoracic ganglion (stellate ganglia). The predicates used to connect these anatomical structures in InterLex are “marksAnatomicalJunction” and “anatomicalJunctionOf” (Table 1). The main difference between these relationships is that marksAnatomicalJunction is used to describe an anatomical point used as a point of reference for an image or scaffold. At the same time, anatomicalJunctionOf refers to the whole surface where the junction occurs.

Custom ontology terms comprise 109 terms that represent non-anatomical terms used when annotating or segmenting data or, more commonly, compositions of anatomical terms. A few examples of these terms are Non-biological empty space (ilx:0738427) or perineurium of vagus nerve (inner edge)(ILX:0739234). The term engineering for these composite parts involves relating them to the appropriate anatomical structures. For instance, the term “ganglia of the inner submucosal nerve plexus”, “is Part Of”, “inner submucosal nerve plexus’.

To bind new terms to the appropriate organs so that they can be returned for an organ-specific query through SciGraph, we use the generic relationship “IncludeForSPARC”. In the above example, urinary bladder-includeForSPARC-dorsal part of serosa of urinary bladder relationship exists a triple within InterLex. This relationship is also used to bind existing terms to an organ when the necessary knowledge engineering is not present in the source ontology to return it for an organ-specific query. For example, the individual spinal nerves were not included in the spinal cord drop-down list accessed via the MBF tools. An investigator requested the terms T3 and C8 spinal nerves, which already existed in the SPARC Vocabulary via FMA, although they are not considered proper parts of the spinal cord. However, in an experimental prep of the spinal cord, it is very likely that the spinal nerves may also be present. The “IncludeForSPARC” relationship was therefore used to link spinal nerves to the spinal cord so they would appear in the drop-down menu.

Through the includeForSPARC relationship, we enhanced the term lists for the multiple organs: peripheral nervous system (75 terms), spinal cord (44 terms), urinary bladder (22 terms), lower urinary tract (28 terms), colon (19 terms), and heart (11 terms). Terms assigned to relationship tag IncludedForSPARC, can be assigned to multiple organ systems to make the term easily findable by the investigator in an external tool or search. This term engineering step allowed for a collection of organ-specific terms to be accessible for annotating images with MBF Bioscience tools (Figure 4). In addition, SPARC Vocabulary term lists per organ can be retrieved as a dynamic query against SciGraph using FMA identifiers for each organ by using the following URL (https://scicrunch.org/api/1/sparc-scigraph/dynamic/prod/sparc/organParts/{FMA:ID}) with the url parameter bringing in the FMA curie identifier. For example, if a user wants to query terms within the SPARC vocabulary mapped to the heart (FMA:7088), the user will use the following URL https://scicrunch.org/api/1/sparc-scigraph/dynamic/prod/sparc/organParts/FMA:7088.

### SAWG term review

Of the 303 anatomical terms added to InterLex, 172 terms underwent review by the SAWG. The remaining terms were either submitted before the SAWG review process was in place or were handled by the SPARC curators without review. Of these 172 terms, 140 were immediately accepted, 30 terms required editing prior to accepting, and 2 terms were rejected. An example of the review process for 3 terms is illustrated in Figure 10:

**Figure 10.**
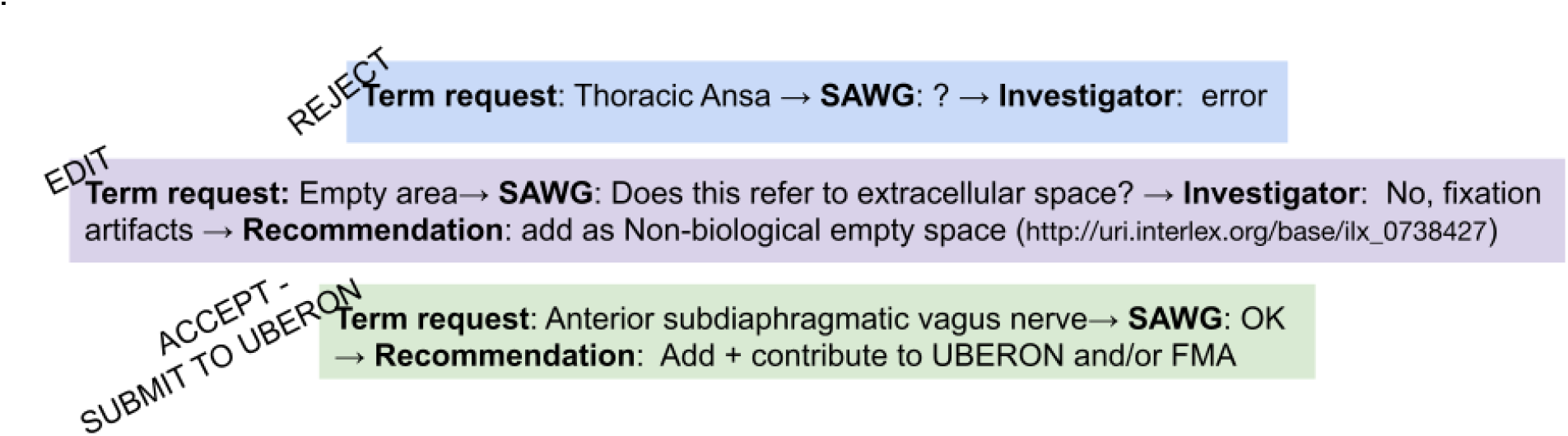
SAWG term review. Examples of the SAWG review process. e.g., The Thoracic ansa was submitted to the pipeline and reviewed by the SAWG since the SAWG did not recognize the term after additional investigation. Once contacted, the investigator indicated it was an error.

#### Rejected term

The term “Thoracic ansa’’ was not recognized by SAWG nor could it be found in the literature. The SAWG requested more information from the investigator, who found the term to be an error. The term was rejected by the SAWG and was not entered into InterLex.

#### Edited term

The term “Empty space” was submitted without a definition and was ambiguous, as it could refer to either the lumen of a structure or a fixation artifact observed in a microscopic image. The investigator clarified that it was the latter, and the SAWG recommended that the term be changed to “Non biological empty space”, which the investigator accepted.

#### Accepted term

The term “Anterior subdiaphragmatic vagus nerve” was determined to be a bona fide anatomical structure; that is, it is referred to in the literature and various parcellation schemes that was missing from the core ontologies. This term was added to the SPARC Vocabulary through the “includeForSPARC” relationship and contributed to UBERON so it would be available to the wider biomedical community.

In total, 8 terms were submitted to UBERON following this review, via workflow specified by the UBERON from the term request pipeline (https://github.com/obophenotype/uberon/blob/sparc-term-request-flow/README-editors.txt). In addition to the 3 terms mentioned previously, 5 additional terms that existed in the species-specific ontologies EMAPA and FMA were recommended for inclusion in UBERON. The terms are in the process of being submitting to UBERON.

## Discussion

This paper introduces the SPARC vocabulary used to annotate data, models, scaffolds, and knowledge within the SPARC program. In addition, it documents a workflow, tools and infrastructure, and review process for working with and adding terms to the vocabulary. This process balances the need for the use of FAIR community ontologies to facilitate integration across databases with the need for deep annotation of experimental data and models of the autonomic nervous system (Osanlouy et al. 2021; Bandrowski et al. 2021). By introducing a curatorial process, including expert review, we ensure that the terms are accurate, non-duplicative (i.e., not already present in an ontology), and more FAIR. We show that by employing this pipeline, we improve the quality of custom terms used for annotation while increasing the degree of mapping of SPARC data to FAIR vocabularies and enriching community ontologies in our domain of expertise.

programs such as SPARC, which generate a large amount of specialized data and tools, face a challenge when trying to implement FAIR vocabularies (Balhoff et al., 2014; Dietze et al., 2014). Community ontologies provide a backbone of semantics for the core entities likely to be encountered and the basic relationships that knit these into a current view of how they are organized. Tools such as BioPortal (Musen et al., 2008), the Open Biological and Biomedical Ontology (OBO) Foundry (Smith et al., 2007), and the Ontology Lookup Service (OLS) maintained by EMBL-EBI (Côté et al., 2006) significantly assist the field by allowing toolmakers and researchers to access and search for ontology terms across ontologies (Balhoff et al., 2014). These tools work reasonably well for dedicated curators working on knowledge bases. However, when it comes to non-dedicated curators employing these vocabularies within real life experimental use cases, there is a need for better tools to search, access, extend and work with these ontologies efficiently and in real-time to serve a particular context. Knowledge engineering is a specialized skill and a time-consuming process, and cannot be done in real-time as curators, investigators, and developers seek to annotate their products in a consistent way (Balhoff et al., 2014).

### Searching for terms

Our experience in SPARC highlights the difficulty of working with large ontologies, as searching across their entirety, including terms and relationships, remains a challenge. Ontologies sit between the realm of human knowledge and computer code (Rzhetsky and Evans, 2011). They contain some classes and relationships that are understandable to a domain expert, but also a lot of opaque relationships and intermediate classes that are difficult for non-experts to understand (Dietze et al., 2014). Nonetheless, for deep annotation of biomedical data, the domain expert must interact with these ontologies to correctly apply them. Thus, a search system or tool must be able to provide the necessary content and context for a domain expert to be able to choose the correct term(s) without overwhelming them.

A large percentage (40%; 120/303) of the terms that were submitted through our term request pipeline already existed in the SPARC Vocabulary, but were not found by users. As discussed earlier, SPARC users can only access the full SPARC Vocabulary, including the entirety of FMA, via our SciGraph instance, which requires computer skills and familiarity with the CYPHER query language. The more user-friendly forms of access-either through searching InterLex or via the MBF annotation tools-provide access only to a subset of the entire SPARC Vocabulary. Thus, to access the full vocabulary through a reasonably user-friendly GUI, annotators would have to search InterLex and then expand to other interfaces such as OLS, OBO Foundry or BioPortal. While dedicated curators might exert this amount of effort, individual investigators and developers do not. Even curators may face a challenge if they are trying to determine how a particular term is related to other terms, as most of the services do not include the entire set of relationships or reasoned hierarchies. FMA poses a particular challenge in this regard, as it is a large complex ontology. OLS and OBO Foundry only contain only a subset of FMA (https://www.ebi.ac.uk/ols/ontologies/fma). The latest version of FMA in Bioportal is 2019 and it is hard to determine whether this is the latest version or not. The University of Washington, which maintains FMA, makes a browsing tool available but it is difficult to use for a non-dedicated user. In the future, ensuring that users can query the entirety of the SPARC Vocabulary through an easy-to-use interface would make the process more efficient.

### Adding and extending ontologies

Almost all ontologies in common use across biomedicine maintain a term tracker that allows a user to suggest a term. However, most community ontologies are under-resourced and cannot deal with requests in real time. Just because a term is requested does not mean it will be added, and the amount of information that may be required to submit the term may be more than the requester is willing to provide. Neither the OBO Foundry or OLS has a centralized term request pipeline; rather they provide links to the individual term request workflows for their ontology of interest. Therefore, to request new terms, investigators must use the term request pipelines for each individual ontology which may be a lengthy process (see (Mungall, 2021), for a detailed essay on adding terms to ontologies). In contrast, Bioportal does provide the means to propose a new term when using a particular ontology, by accessing the “Create a proposal” function under the Notes field, although a non-dedicated domain expert may have difficulty finding and/or understanding its use. It also provides a more generic provisional term request via an API that will assign a temporary identifier to a proposed term that does not have to be submitted to a particular ontology, although the submitter can recommend one (https://ncbo.bioontology.org/wiki/BioPortal_Provisional_Terms). The SPARC term request pipeline provides 2 simple interfaces for requesting new terms, InterLex and a Google Sheet. InterLex allows terms to be added directly through a readily accessible form-based graphical user interface (GUI), assigns them an identifier, and makes them immediately available for annotation. If terms aren’t needed right away or if there are too many to add manually, users may submit via a Google Spreadsheet. In this way, users with minimal ontology expertise can contribute terminology to SPARC.

The development of the term request pipeline was informed by our efforts in developing vocabularies for neuroscience through the Neuroscience Information Framework (NIF) project, an NIH-Blueprint-funded project to develop a resource description framework for neuroscience resources (Gardner et al., 2008). NIF developed a set of ontologies, the NIFSTD, that focused on coverage in the major domains of neuroscience, including anatomy, cells, subcellular structures, etc. (Bug et al., 2008). A goal of NIF was to make it easy for those who wanted to contribute ontology expertise to NIFSTD to be able to add, edit and review terms. Towards this aim, the NIF project launched the NeuroLex wiki in 2008 (Larson and Martone, 2013), a semantic wiki built on the Semantic MediaWiki platform with specialized extensions that allowed someone with minimal experience working with ontologies to add, modify and link ontology terms. It was initially seeded with the NIFSTD and was a fairly successful platform, tracking over 25,000 terms. It lowered the barrier to entry for using formal vocabularies and ontologies by allowing domain experts with limited knowledge of ontology engineering to provide a view of how these terms linked to others (Hamilton et al., 2012). Over the period of 2009-2013, it received more than 200,000 edits by > 100 different users (Larson and Martone, 2013).

Neurolex was discontinued in 2018, as the customized extensions to the Semantic MediaWiki platform became increasingly difficult to support. InterLex was developed as a replacement and all content from Neurolex was ported to the new platform. Unlike Neurolex, as InterLex was developed after the FAIR principles became well known, it issues FAIR identifiers in the form of CURIES as well as full URIs. InterLex is being used as a vocabulary management system by several neuroscience-related programs in addition to SPARC, including ReproNIM (Kennedy et al., 2019) and the Open Data Commons for Spinal Cord Injury (Callahan et al., 2017). These latter two projects are also using InterLex to store common data elements and custom data elements and relate them to ontological terms.

Neurolex and InterLex both take a bottom-up approach to the creation of formal vocabularies and ontologies for neuroscience by creating a readily accessible, community-driven lexicon of useful terms. These terms can be used on their own, grouped into term sets through simple tagging and relationships, or subjected to more elaborate term engineering and imported into ontologies like NIFSTD (Imam et al., 2012) or the SPARC Vocabularies as needed. As with our current pipeline, it has always been NIF’s practice to contribute back terms to enrich community ontologies with neuroscience content where appropriate. NIF contributed a significant number of subcellular terms to the Gene Ontology Cell Component ontology (Roncaglia et al., 2013) and added neuroanatomical content to RadLex and NeuroNames (Turner et al., 2010). It is interesting that of the 300 or so terms contributed to SPARC, only eight terms were considered to be good candidates for UBERON, suggesting that coverage of the ANS and related structures in UBERON is very good. The majority of missing terms requested by SPARC are not the types of atomic entities usually included in ontologies, but represent often ad hoc compositions of existing terms required in an experimental context (Dietze et al., 2014). As there are an astronomical number of these compositional terms, ontologies typically do not precompose them, requiring the user to perform the necessary knowledge engineering to create the necessary linkages between existing classes. Tools like TermGenie (Dietze et al., 2014) provide templates which assist in this process, but we chose to implement the engineering by defining a pattern and having semi-expert curators perform the necessary knowledge engineering to relate these terms to their component structures and any necessary qualifiers. The SPARC term request pipeline, therefore, increases the FAIRness of SPARC data by ensuring these more granular annotations, which traditionally would have been tagged with free text, are mapped to their component structures.

Similarly, many terms that were submitted via the MBF pipeline represented terms that were in community ontologies but were missing from the organ-specific term lists accessed by these investigators. We assembled the organ specific term lists using SPARQL queries against UBERON and FMA that traverse the partonomy chain associated with a particular organ. Many of the terms requested, however, belong to other organs that are found in proximity to this structure in experimental preparations but don’t properly belong to the parent organ. In this case, we appended the contributed terms to the main list through the includeForSPARC relationship so that they became available to future annotators. In that way, the provenance is clear for anyone using the SPARC Vocabulary that these relationships come via SPARC.

### Quality control

Although InterLex makes it easier for anyone to add terms and work with ontologies, as one might imagine, the quality of the term metadata and relationships is highly variable. Before the term request pipeline and curatorial review was established, multiple terms were added with no definitions or relationships provided, limiting their capacity for reuse. In order to ensure that SPARC is being annotated with high quality terminologies, we therefore instituted a curatorial and expert review process specifically for any anatomical terminology contributed. Only curated terms are incorporated into the SPARC Vocabulary available through SciGraph. However, all contributed terms remain in InterLex, unless they were rejected by the SAWG as erroneous and therefore not added to or removed from InterLex. Since instituting this process, most terms submitted have definitions and provenance, and additional relationships provided, indicating that building high quality vocabularies on a community platform benefits from curatorial oversight and quality checks (Balhoff et al., 2014).

### Future directions

Although the basic infrastructure is in place for managing term additions and basic curation, several major improvements are planned to streamline and manage the process. A top priority is to implement a fully-configured third party curatorial workflow on top of the terms submitted to InterLex. This system will include an interactive dashboard where a curator can monitor submitted terms and perform basic curatorial functions such as tagging. It will include a notification system that notifies both curators and submitters when terms are added, edited, undergone SAWG review, deprecated or annotated. While InterLex has the basic pieces of these functions, a dedicated curatorial interface will greatly enhance and help manage the workflow.

A second priority is a tighter integration with external ontology services. The goal is for InterLex to establish a push-pull relationship with community ontologies, allowing us to pull terms directly from a service like BioPortal, OLS or Ontobee, and in turn, notify any source ontologies of new terms within their domains. Ideally, if the changes are accepted, they would be automatically pushed back to InterLex. Such a service would make it much easier to keep InterLex in sync with community ontologies. Towards this end, we are implementing the ability to “follow” a particular branch of InterLex so that 3rd parties can be notified if terms are added or edited within a particular branch.

Finally, the way InterLex handles versions is being rethought. Currently, while a user is able to view previous versions of a term within InterLex via its history, the URI only points to the latest version of a term and InterLex does not define stable URIs for interim versions. Given that the vocabularies in InterLex can be fluid, one should be able to point to a specific version in a given vocabulary as some types of edits may alter the definition or relationship of a term.

Other planned improvements are a bulk upload feature for the submission of multiple terms and making it easier for a research group to assemble, display and download a customized term set for their program.

## Conclusions

The SPARC vocabulary and infrastructure assist researchers and developers in creating and using FAIR vocabularies by providing the flexibility for easy addition of new terms that adhere to FAIR principles. InterLex serves as a workspace for viewing and working with vocabularies that are accessible to those with limited expertise in knowledge engineering. When coupled to a curatorial service, the necessary knowledge engineering can be performed to link these terms to existing community ontologies. Using this infrastructure, we have shown that we increase the “FAIRness” of SPARC data by extending the concept of FAIR to terms that may fall through the cracks of current ontologies. We conclude that the term request pipeline serves as a useful adjunct to community ontologies for annotating experimental data in support of FAIR.

## Acknowledgments

This work was supported by NIH grant 3OT2OD030541 from the Office of the Director through the Stimulating Peripheral Activity to Relieve Conditions (SPARC) program. We thank Drs. Anita Bandrowski and Jyl Boline for their helpful comments.

## Data Availability Statement

Publicly available term history from this study is available at https://scicrunch.org/sawg/interlex/dashboard-history?origCid=504&page=1 and https://doi.org/10.5281/zenodo.5703661.

## Contribution to the Field Statement

This manuscript describes a flexible system for building and annotating with FAIR vocabularies developed for the Stimulating Peripheral Activity to Relieve Conditions (SPARC) program, a program to accelerate the development of treatments for autonomic nervous system (ANS)-based disorders through bioelectronic medicine. We describe an infrastructure for housing, accessing and extending the community ontologies used to annotate data within SPARC, with a focus on neuroanatomical structures. These functions are mediated through Interlex, an on-line vocabulary management system, developed initially by the Neuroscience Information Framework. Terms are added by researchers, knowledge engineers and developers, to annotate SPARC multimodal experimental data, models and knowledge about ANS connectivity. Each added term receives a full URI, metadata and may be connected to other terms through formal relationships. In order to ensure that anatomical terms are of high quality and clearly defined, a term review process was established for anatomical experts to review these terms. We provide a solution to the problem of annotating experimental data, which often requires more granular terms than are provided by community ontologies, and show by incorporating both a term request pipeline and infrastructure increases the FAIRness of SPARC products.

